# Global analysis of specificity determinants in eukaryotic protein kinases

**DOI:** 10.1101/195115

**Authors:** David Bradley, Cristina Viéitez, Vinothini Rajeeve, Pedro R. Cutillas, Pedro Beltrao

## Abstract

Protein kinases lie at the heart of cell signalling processes, constitute one of the largest human domain families and are often mutated in disease. Kinase target recognition at the active site is in part determined by a few amino acids around the phosphoacceptor residue. These preferences vary across kinases and despite the increased knowledge of target substrates little is known about how most preferences are encoded in the kinase sequence and how these preferences evolve. Here, we used alignment-based approaches to identify 30 putative specificity determinant residues (SDRs) for 16 preferences. These were studied using structural models and were validated by activity assays of mutant kinases. Mutation data from patient cancer samples revealed that kinase specificity is often targeted in cancer to a greater extent than catalytic residues. Throughout evolution we observed that kinase specificity is strongly conserved across orthologs but can diverge after gene duplication as illustrated by the evolution of the G-protein coupled receptor kinase family. The identified SDRs can be used to predict kinase specificity from sequence and aid in the interpretation of evolutionary or disease-related genomic variants.

## Introduction

Protein post-translational modifications (PTMs) constitute one of the fastest mechanisms of control of protein function and protein phosphorylation is the most extensive and well characterized PTM. Protein kinases catalyse the phosphorylation of their target substrates, including other kinases, working in complex signalling networks that are capable of information processing and decision making. These signalling networks are involved in almost all cellular processes and mutations in protein kinases are often associated with disease (Lahiry et al. 2010; Stenberg, Riikonen, and Vihinen 2000; Brognard and Hunter 2011). In addition, cross-species studies have shown that protein phosphorylation and kinase-substrate interactions can diverge at a very fast pace, suggesting that changes in post-translational control can be a driver of phenotypic diversity (Beltrao et al. 2009; Freschi, Osseni, and Landry 2014; Studer et al. 2016). Understanding kinase signalling networks remains a difficult challenge, in particular because only a small fraction of the known phosphorylation sites can be assigned to their effector kinases.

There are 518 known human protein kinases (Manning et al. 2002), and their specificity of substrate recognition is shaped by the structural and chemical characteristics of both kinase and substrate (Ubersax and Ferrell 2007). The general fold of different kinases is quite similar and the specificity of kinases is, in part, determined by changes near the binding pocket. Kinases are thought to recognise a contiguous motif around the phosphosite (four/five amino acids on either side of the P-site) (Knighton et al. 1991; Pearson and Kemp 1991; Pinna and Ruzzene 1996; Amanchy et al. 2007) usually termed the kinase target motif. These target motif preferences are most often very degenerate with only a small number of key residues strongly contributing to the recognition. While these sequence preferences are thought to be important for target recognition, additional mechanisms contribute to specificity including: docking motifs; interaction with protein scaffolds; co-expression and co-localization (Biondi and Nebreda 2003; Holland and Cooper 1999). Sequence analysis has identified 9 kinase groups (AGC, CAMK, CMGC, RGC, TK, TKL, STE, CKI and other) but only a few kinase groups have clear differences in target preferences that are shared with most members of the group. For example the CMGC kinases tend to phosphorylate serine and threonine residues that have proline at position +1 relative to the phospho-acceptor (Kannan and Neuwald 2004). However, for most kinase groups the preferences for residues around the target phospho-acceptor cannot be easily predicted from the primary sequence.

In previous studies of kinase specificity, the analysis of protein structures (Brinkworth, Breinl, and Kobe 2003; Saunders et al. 2008) and machine learning methods (Creixell, Palmeri, et al. 2015) have been used to identify positions within the kinase domain that determine kinase specificity – so called specificity determinant residues (SDRs). However, these approaches do not attempt to study the structural basis by which specific target preferences are determined. Methods based on protein kinase alignments can achieve this, but have only been used to study a few kinase groups so far (Kannan et al. 2007; Kannan and Neuwald 2004), or have been restricted to a single model organism (Mok et al. 2010). Here we have used alignment and structure based methods to identify and rationalise determinants of kinase specificity. We have identified SDRs for 16 target site preferences and show that these can be used to accurately predict kinase specificity. We provide detailed structural characterizations for many determinants and study how these are mutated in cancer or during evolution. We show how the knowledge of SDRs can be combined with ancestral sequence reconstructions to study the evolution of kinase specificity using as an example the G-protein coupled receptor kinase family.

## Results

### Identification of kinase specificity-determining residues and modelling of the kinase-substrate interface

To study kinase target preferences we compiled a list of 9005 experimentally validated and unique kinase-phosphosite relations for human, mouse and yeast kinases. Protein kinase specificities were modelled in the form of position probability matrices (PPMs) for 179 kinases, representing a fraction of the kinome of these species (human: 126/478, mouse: 35/504, *S. cerevisiae*: 18/116). For further analysis, we selected 135 high-confidence PPMs (87 human, 30 mouse, 18 yeast) that could successfully discriminate between target and non-target phosphorylation sites (see Methods). For serine/threonine kinases, consistent evidence of active site selectivity is broadly apparent for the -3 and +1 positions relative to the phosphoacceptor, and to a lesser extent the -2 position (**Figure 1a**). These constraints correspond mainly to the well-established preferences for basic side chains (arginine or lysine) at the -3 and/or -2 position, and in most CMGC kinases for proline at the +1 position. Tyrosine kinases however show little evidence of strong substrate preference at the active site, and were excluded from further analysis as there were too few high-quality PPMs (16) for the reliable detection of tyrosine kinase SDRs. These trends only describe the most common modes of recognition shared across many kinases, and individual kinases can show preference for positions beyond these sites. All 135 high confidence kinase specificity models are summarized in **Supplementary Table 1**.

**Figure 1.**
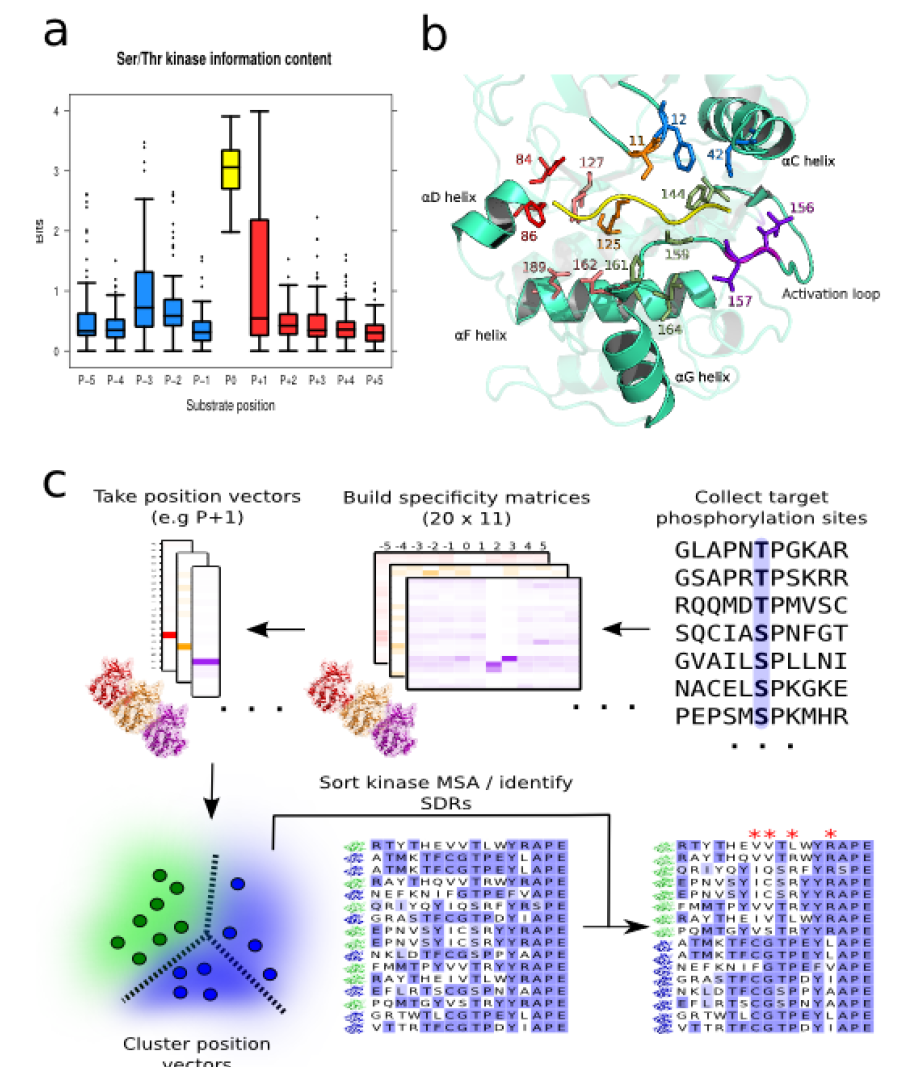
Features of kinase target interaction and pipeline for SDR identification. a) Sequence constraint for substrate positions -5 to +5 for 119 serine/threonine kinases, measured as the bit value for the corresponding column of the kinase PSSM. b) Interface between a protein kinase (human protein kinase A) and substrate peptide at the substrate-binding site. Kinase residues that commonly bind the substrate peptide (yellow) are represented in stick format and coloured according to the corresponding substrate position (-3: red, -2: pink, -1: orange, +1: green, +2: blue, +3: purple). Residue numbering represents the relevant positions of the Pfam protein kinase domain (PF00069) c) Semi-automated pipeline for the inference of putative kinase SDRs (specificity-determining residues). The first step involves the construction of many kinase PPMs from known target phosphorylation sites. Vectors corresponding to a substrate position of interest (e.g. +1) are then retrieved from each PPM. An unsupervised learning approach (i.e. clustering) then identifies kinases with a common position-based preference (e.g. for proline at +1). Alignment positions that best discriminate kinases belonging to one cluster from all others are then identified using automated tools for SDR detection.

With this information, we then attempted to understand more broadly the relationship between protein kinases and substrates at the active site by employing structural models (**Figure 1b**) and kinase sequence alignments (**Figure 1c**). We compiled 12 serine/threonine non-redundant experimental structural models of kinases in complex with substrates, in addition to 4 serine/threonine autophosphorylation complexes (Xu et al. 2015) (see full list in **Supplementary Table 2**). Kinase-substrate homology models for kinases of interest not represented in this compilation of experimental models were also generated. A structural profile of substrate binding from position -5 to position +4 is given in **Supplementary Figure 1**. The kinase positions most frequently in contact with the target peptide are highlighted also in **Figure 1b**. When referring to specific amino acids in the kinase, the single-letter code is used followed by the position of the residue based on the Pfam protein kinase domain model (PF00069).

We developed a sequence alignment-based protocol for the automated detection of putative specificity-determining residues (**Methods, Figure 1c**). Briefly, the target preferences described as PPMs were clustered to identify groups of kinases with shared preferences at a position of interest. Putative SDRs are then inferred as those residues that discriminate kinases with the common substrate preference (e.g. proline at the +1 position or P+1) from other kinases (**Figure 1c**). Using this approach we identified 30 predicted SDRs for 16 preferences (**Figure 2a**) found across the sequence/structure of the kinase domain (**Figure 2b**). Not surprisingly SDRs tend to cluster near the binding pocket (**Figure 2c)** with 33% near the substrate compared to ∼12% for any kinase position (Fisher p < 0.01). To assess the accuracy of these SDRs we tested if these could be used to predict the specificity of kinases from their sequence alone. For this we built sequence-based classifiers for the five preferences supported by at least 20 positive examples in the study dataset – P+1, P-2, R-2, R-3, and L-5. We used a cross-validation procedure where kinase sequences left out from the model training were later used for testing (**see Methods**). These models showed very strong performance with respective cross-validation AUC values of between 0.82 and 0.99 (**Supplementary Figure 2**). This shows that, for these 5 specificities, the determinant residues can correctly predict the specificity of unseen kinases from their sequence alone, suggesting that the SDRs we have identified are broadly accurate.

**Figure 2.**
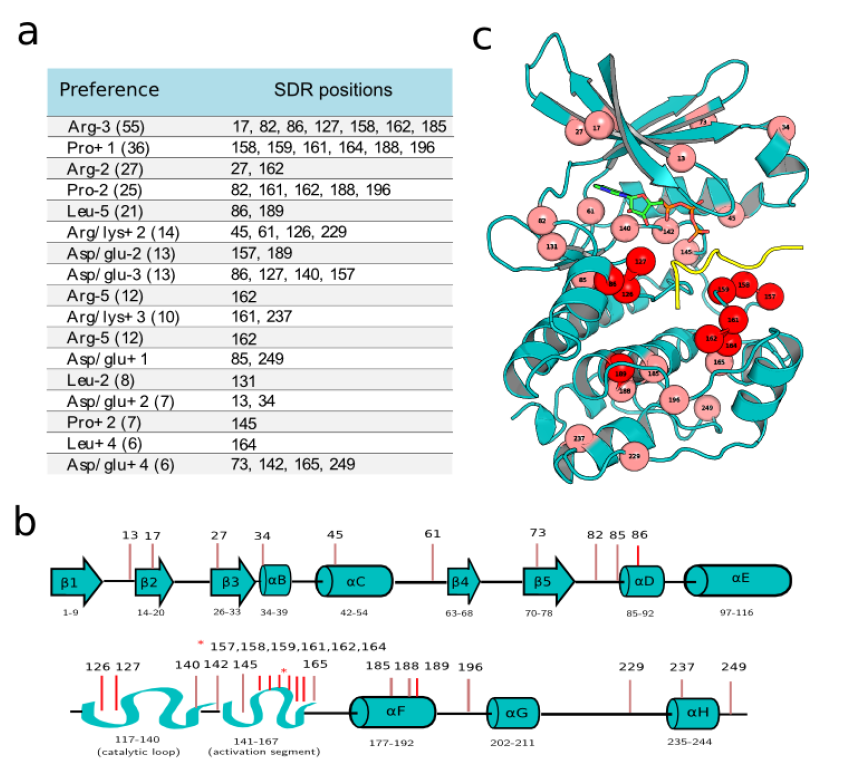
Position of identified SDRs along the kinase sequence and structure. All putative kinase SDRs from our analysis are a) listed in a table with their corresponding position preferences b) mapped to a 1D representation of the kinase secondary structure c) mapped to a kinase-substrate complex structure (PDB: 1atp). The SDRs colored in dark red b) and c) represent positions within 4 Angstroms of the substrate peptide. Residue numbering represents the relevant positions of the Pfam protein kinase domain (PF00069)

### Structural characterization of kinase SDRs

Most of the predicted SDRs have not been described before and can be further studied by analysis of structural models. We have used available co-crystal co-ordinates where possible and models of relevant kinase-substrate complexes were alternatively generated using empirical complexes as a template (**see Methods**). Using these models we could suggest a structural rationale for SDRs of 8 target site preferences that are detailed in **Supplementary Figure 3**. These include the preferences for arginine at positions -3 and - 2; proline at positions -2 and +1; leucine at positions +4 and -5 and for aspartate/glutamate at position +1 for AGC and CMGC kinases. Some of the SDRs have been previously identified in other studies underscoring the validity of our approach. For example, four of the six putative SDRs identified here for the proline +1 preference map to the kinase +1 binding pocket (**Supplementary Figure 3**) and match previously described determinants (Kannan and Neuwald 2004).

**Figure 3.**
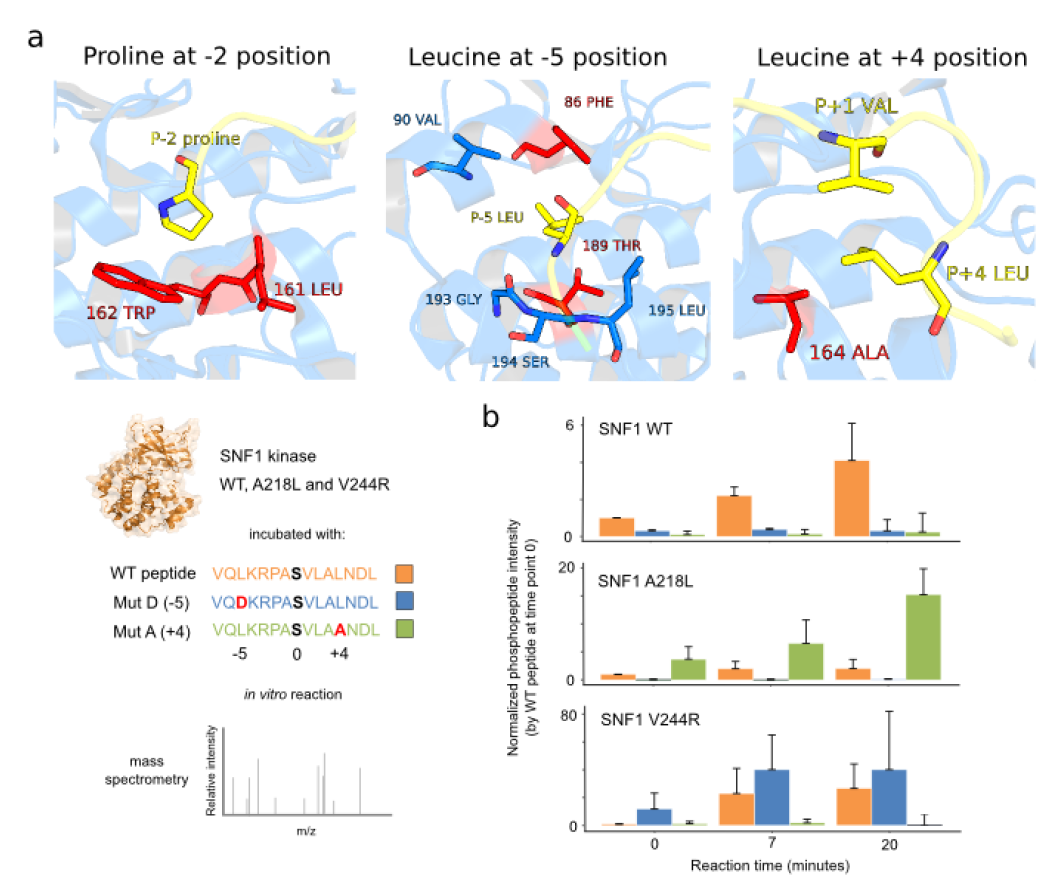
Structural rationale for kinase SDRs and validation experiments. a) Kinase-substrate interface for: proline at position -2 (PDB: 2wo6), leucine at position -5 (PDB: 3iec) and leucine at position +4 (PDB: 3iec). The substrate peptides are colored in yellow, and putative SDRs in red. A structural rationalisation for each preference is provided briefly in the main text ‘structural characterization of kinase SDRs’, and in more detail in **Supplementary Figure 3**. b) Kinase activity assays for SNF1 WT and two mutant versions A218L (the 164 kinase position, an L+4 SDR) and V244R (the 189 kinase position, an L-5 SDR). The 3 kinases were incubated separately with a known SNF1 target peptide with L at +4 and -5 (orange) as well as the mutant versions A+5 (green) and D-5 (blue). Replicates of in vitro reactions were quenched at 0, 7 and 20 mins and the amount of phosphorylation was measured by mass spectrometry. For each kinase and time points the phosphopeptide intensity relative to the WT peptide at time point zero was calculated and the median and standard deviation of 3 biological replicates are plotted.

We highlight in **Figure 3a** SDRs for 3 preferences that are less well studied: proline at position -2 (P-2) and leucine at positions +4 (L+4) and -5 (L-5). There are 25 kinases with a modest P-2 preference including MAPK1, CDK2, and DYRK1A. We identified 5 positions that are putative SDRs for P-2, two of which (161 and 162) are proximal to the residue in interaction models. In position 162, P-2 kinases usually contain a bulky hydrophobic residue (Y or W) not usually found in non-proline-directed kinases (**Supplementary Figure 3**). Both residues at these positions appear to form hydrophobic contacts with P-2 (**Figure 3a**). The domain position 161 was also implicated in the preference for the P+1 specificity mentioned above. The three other putative determinants – 82, 188, and 196 – are unlikely to be direct determinants given their distal position in the protein structure, although we note that 196 was implicated in a previous alignment-based study (Mok et al. 2010). These distal positions may influence the kinase preference through more complex mechanisms such as affecting the dynamics or conformation of the kinase.

We identified 21 kinases (14 CAMK; 5 AGC; 1 CMGC; 1 PRK) with a moderate L-5 preference. Positions 86 and 189 were predicted as determinants where L-5 kinases are marked by hydrophobic amino acids at position 86 and the absence of glutamate at 189. These residues can be observed to line the hydrophobic -5 position pocket of the MARK2 kinase (**Figure 3a**). Position 189 was also recently predicted to be an L-5 determinant from a comparative structural analysis of L-5 and R-5 kinases (Catherine Chen et al. 2017). For the leucine preference at the +4 position we identified six kinases – MARK2, CAMK1, PRKAA1, PRKAA2 (human), PRKAA1 (mouse), and Snf1 (yeast) – and the domain position 164 as the sole putative SDR. This residue is an alanine in five of the kinases listed above (valine in CAMK1). In the MARK2 cocrystal structure, the substrate peptide forms a turn at the +2 position so that the +4 hydrophobic side chain projects towards the kinase pocket of the +1 position and stacks against the +1 residue (**Figure 3a**). The substitution for alanine in place of residues with aliphatic side chains at position 164 in these kinases therefore seems to generate a small binding pocket that allows the L+4 to functionally substitute for the kinase position 164 by stacking against the +1 position.

We have selected two of the above described SDRs for experimental characterization (L-5 and L+4). To test these SDRs we performed in vitro kinase activity assays for SNF1 WT and two mutant versions of the kinase: A218L (the 164 kinase position, an L+4 SDR) and V244R (the 189 kinase position, an L-5 SDR). These 3 kinases were expressed and purified from yeast cells and individually incubated with a SNF1 target peptide of 15 amino acids that contains leucine at +4 and -5 as well as mutant versions with A+4 or D-5. The in vitro kinase reactions were quenched at 0, 7 and 20 minutes and the amount of phosphorylation was measured by mass spectrometry (**Figure 3b**). As predicted the A218L SNF1 showed an increased preference for the A+4 peptide but not for the D-5. The reverse was observed for the V244R SNF1 mutant.

The identification of previously known SDRs, the structural rationale for several of the novel SDRs and the experimental validation of two SDRs, further suggests that we have here identified positions that are crucial for the recognition of kinases with specific preferences. The SDRs identified here can therefore be used to infer the specificity of other kinases from sequence and, as we show below, to study the consequences of mutations within the kinase domain.

### Specificity determinant residues are often mutated in cancer

Some kinase SDRs have been observed to be mutated in cancer and congenital diseases (Creixell, Schoof, et al. 2015; Berthon, Szarek, and Stratakis 2015). Using mutation data from tumour patient samples from TCGA (http://cancergenome.nih.gov/), we have tested for the enrichment of tumour mutations in four categories of kinase residues: catalytic, regulatory, SDR (proximal to substrate), and ‘other’ (**Figure 4a**). SDR residues close to the substrate show a significant enrichment of mutations relative to ‘other’ residues in the kinase domain (Mann-Whitney, p = 0.0006, **Figure 4b**). This enrichment is greater than that observed for catalytic and regulatory sites, highlighting their functional relevance.

**Figure 4.**
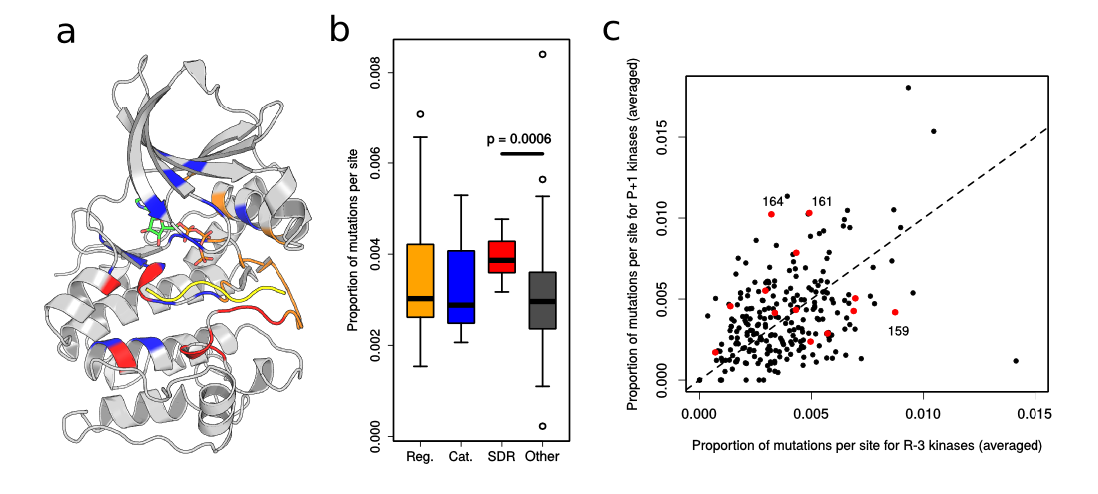
Mutation of SDRs in cancer. a) Kinase domain positions are colored according to their functional category (regulatory: orange, catalytic: blue, SDR: red, ‘other’: grey). The substrate peptide is represented in yellow and ATP in green, orange, and red. b) The fraction of mutations mapping to a given site for a given Ser/Thr kinase were calculated, and then averaged across all Ser/Thr kinases. The different sites are grouped according to their functional category. c) For a given site, the frequency of mutations in arginine-3 kinases (x-axis) and proline+1 kinases (y-axis) is plotted. Putative SDRs are colored in red.

We next sought to determine if the frequency of SDR mutations differs between kinases depending upon their specificity. Given that the specificity models only cover ∼25% of all kinases we used the SDRs of the 5 most common preferences - P+1, P-2, R-2, R-3, and L-5 - to train sequence based predictors of kinase specificity as described above. Using these models we annotated all human kinases having a high probability for one of these specificities (**Supplementary Table 3**). We then compared the frequency of mutations per position for different kinase specificities and found significant differences in the relative mutation frequencies for the P+1 and R-3 positions (represented in **Figure 4c**). For positions 164 and 161 of the +1 position loop exhibit high levels of differential mutation in the proline-directed kinases. For position 161, the MAP kinases in particular are recurrently mutated in independent samples (MAPK1: 3, MAPK8: 3, MAPK11: 2, MAPK1: 1). This position is known to bind to the phosphotyrosine at 157 that exists in MAPKs (Varjosalo et al. 2013). For the predicted R-3 kinases, the glycine 159 residue of the +1 position pocket is found to be commonly mutated, although this relates not to R-3 specificity *per se* but for +1 position binding of most non-CMGC kinases (Zhu et al. 2005). Residues 159 and 164 in particular are critical for specificity and highly conserved within the kinase subgroups, such that mutation to any other amino acid would be expected to abrogate P+1 binding. These results suggest that there is a significant recurrence of cancer mutations targeting kinase specificity and not just kinase activity.

The work above illustrates how knowledge of the SDR residues is useful in understanding the functional consequences of cancer mutations. We next studied the changes in SDR residues during the evolution of protein kinases.

### Divergence of kinase specificity between orthologs

The full extent to which kinase specificity differs between orthologs is not known (C. J. Miller and Turk 2018; Ochoa, Bradley, and Beltrao 2018). To study this we first compared 20 ortholog groups with 65 pairs between a human/mouse and a yeast kinase with experimentally determined specificity. Specificity logos for 3 different examples are given in **Figure 5a** indicating that these tend to be similar. We find that the difference in specificity between orthologs (as calculated by the distance between PPMs) is generally similar to the expected for biological replicates of the same kinase (p = 0.097, Mann-Whitney, two-tailed, **Figure 5b**), but is less than that observed for random human-yeast kinase pairs (p << 0.01, Mann-Whitney, one-tailed, **Figure 5b**). Only 6/65 (9%) of orthologous pairs (including for example the yeast kinases Cmk1/Cmk2, Sky1, and Pkc1) are more divergent than the median distance of random human-yeast kinase pairs. Kinase specificities are therefore highly conserved in general between human/mouse and *S. cerevisiae* even though they diverged more than 1000 million years ago (Doolittle et al. 1996).

**Figure 5.**
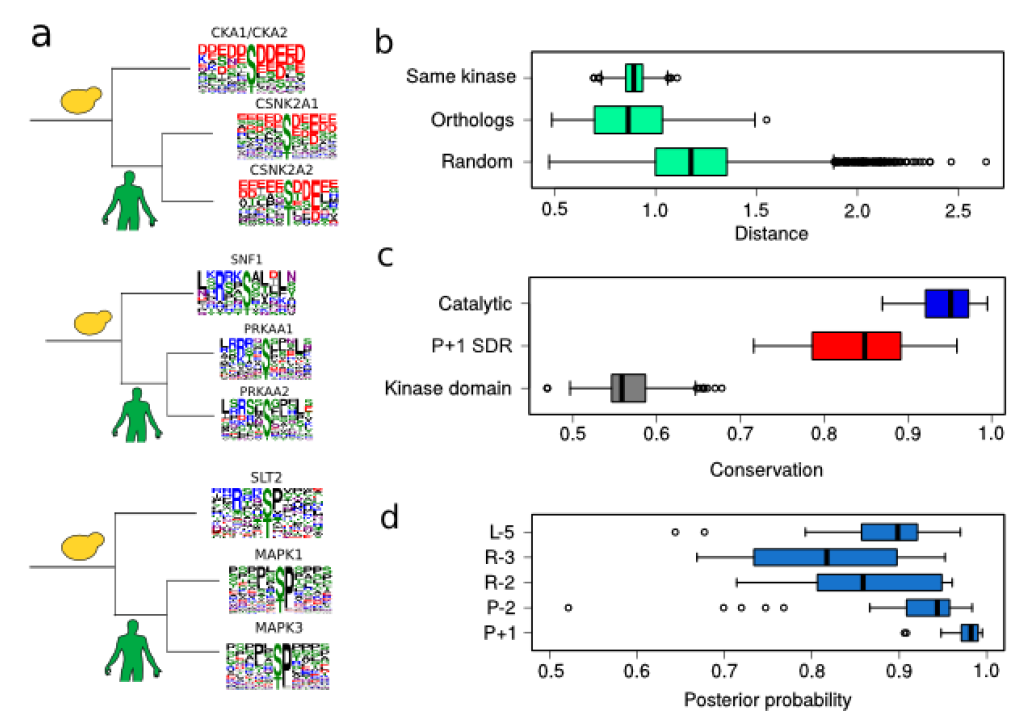
– Evolution of specificity for orthologous kinases. a) Human and yeast kinase specificity logos for three different orthologous groups b) Distribution of matrix distances between PPMs generated from phosphosite subsamples of the same kinase (top), orthologous yeast and human/mouse pairs (middle) and random human-yeast pairs (bottom) c) Conservation of domain residues, SDRs, and catalytic residues for the proline+1 specificity. Each data point represents the average conservation (among kinase domain positions, SDR, or catalytic residues) for an alignment of orthologous kinases where the human kinase is a predicted proline+1 kinase. d) Conservation of specificity for kinases orthologous to human kinases of predicted specificity (L-5, R-3, R-2, P-2, P+1). Each data point represents the average posterior probability (across all kinases in an orthologous group) that the specificity has been conserved.

We next used the identified SDRs to investigate the divergence of specificity between orthologs. We focused our analysis on the 5 specificities we can reliably predict from sequence as described above: P+1, P-2, R-2, R-3, and L-5. Orthologs were retrieved from the Ensembl Genomes Compara database (1210 species) for each human kinase predicted to have one of the five specificities (i.e for P+1, P-2, R-2, R-3, or L-5). SDRs for each of the five specificities show a much higher sequence conservation than other kinase residues, although lower than observed for the essential catalytic residues (**Figure 5c**, **Supplementary Figure 4**). Predictions of ortholog specificity however suggest that this modest sequence variation among SDRs rarely alters kinase specificity (**Figure 5d**). Specifically, we predict divergence (posterior probability < 0.5) for only 5% of orthologous groups. In one of the few examples, the Wee2 protein in human features a hydrophobic -5 binding pocket, but this is the case for vertebrate sequences only. For the 5 specificity classes and for *Arabidopsis thaliana* orthologs of human kinases, we predict that the ortholog specificity has diverged in only 12% of cases.

Taken together, these results demonstrate that kinase specificities tend to be highly conserved across orthologs even between species separated by 1 million years of evolution.

### Divergence of kinase specificity within the GRK family

We then selected the GRK (G-protein coupled receptor kinase) kinase family as specific detailed case study of the evolution of target specificity. The GRK family is one of 15 families belonging to the AGC group (**Figure 6a**) (Manning et al. 2002). However, they have diverged from the characteristic basic residue preferences at positions -2/-5 and -3 of the AGC group (Lodowski et al. 2006). GRK2 for example is specific for aspartate/glutamate at position -3 (Onorato et al. 1991; Lodowski et al. 2006), and in the GRK5 model presented here the R-3 signature is absent (**Figure 6b**). The GRK family is divided into the BARK (β-adrenergic receptor kinase) subfamily – comprising GRK2 (ADRBK1) and GRK3 (ADRBK2) in human – and the GRK subfamily – comprising GRK1 (rhodopsin kinase), GRK4, GRK5, GRK6, and GRK7 (Manning et al. 2002). We have taken a taxonomically broad sample of 163 GRK kinase sequences to generate a comprehensive phylogeny (**Figure 6a**, Methods). From this, a maximum-likelihood reconstruction of ancestral sequence states has been performed (Methods) in order to study the evolution of substrate preferences on the basis of our detailed understanding of kinase SDRs.

**Figure 6.**
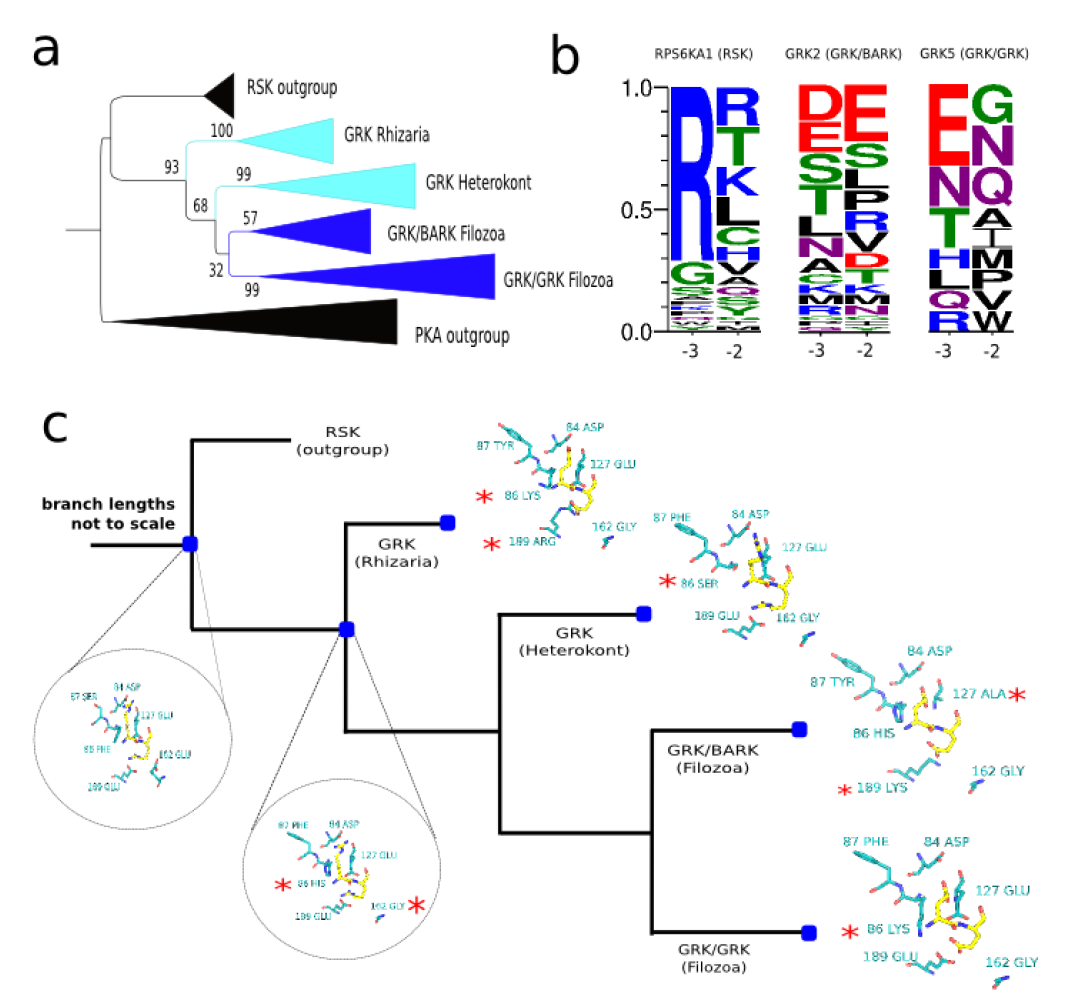
Evolution of GRK family specificity. a) Phylogeny of kinases in the GRK family, including an outgroup of RSK kinases in human. The supporting number of bootstrap replicates (/100) for relevant clades and bifurcations is represented. b) Logos at positions -3 and -2 for human RSPS6KA1 (RSK kinase), human GRK2 (GRK/BARK kinase), and human GRK5 (GRK/GRK kinase). Sequence logos were generated from target phosphorylation sites. c) Representation of substrate position -2 and -3 (yellow), and their corresponding kinase binding pockets (cyan) for extant kinases and predicted ancestral sequences. Substitutions in the binding pocket are denoted by a red asterisk.

The topology of the tree is in general agreement with a previously published GRK phylogeny (Mushegian, Gurevich, and Gurevich 2012). Focusing on the specificity at the - 2 and -3 positions (**Figure 6c and Supplementary Figure 5**), two substitutions between the ancestor of RSK and GRK kinases and the ancestor of all GRK kinases likely caused a reduced preference for arginine at -3 and -2 positions. The substitution of glutamate for glycine at position 162 – an R-3 and R-2 determinant (**Supplementary Figure 3**) – and the substitution of phenylalanine at position 86, most likely either to histidine or to lysine. From this ancestral node towards the Rhizarian lineage an additional substitution of glutamate at 189 for arginine likely drove the complete switch from R-2/R-3 to a novel aspartate/glutamate preference at the -2 position. This 86K/189R pair is analogous to the 127E/189E pair found in basophilic kinases. In the heterokont lineages, the histidine/lysine at position 86 in the ancestor of GRK kinases was substituted for serine and while these kinases retained the 86E/189E pair, the R-2 and R-3 specificities are likely to be attenuated or eliminated given the substitutions at positions 86 and 162. The BARK kinases had two charge altering substitutions – E127A and E189K – that likely generated the preference for aspartate/glutamate at the -2 and -3 positions as observed in extant GRK2 kinases (**Figure 6b**). Finally, in the GRK subfamily, a lysine residue (or arginine in GRK1) is usually found at position 86. Notably, no R-2/R-3/R-5 preference is evident for GRK5 (**Figure 6b**), suggesting that the described substitutions (E162G and F86K) were sufficient to eliminate this specificity.

The GRK family illustrates how the target preference of a kinase can change after kinase duplication via the substitution of a few key residues. It also illustrates one example where distantly-related kinase orthologs may have diverged when comparing the metazoa GRKs to their rhizaria homologs that diverged around 1700 million years ago (Kumar et al. 2017).

## Discussion

We have here addressed the challenge of identifying which residues determine kinase preferences towards specific amino-acids at specific positions around the target phosphosite. Initial studies of kinase determinants used structures of kinases in complex with target peptides to identify SDR residues as those important for substrate binding (Brinkworth, Breinl, and Kobe 2003; Zhu et al. 2005). A more recent work has used a machine learning approach to identify SDR residues as those that globally maximized the specificity predictive power (Creixell, Palmeri, et al. 2015). These approaches have identified SDR positions but do not assign positions and residues according to specific target preferences (e.g. R-3 or P+1). Alternatively, alignment-based approaches can be used to identify residues that contribute to particular preferences but so far have been restricted to one kinase group at a time (Kannan et al. 2007; Kannan and Neuwald 2004), or a single model organism (Mok et al. 2010). Here we have combined a statistical analysis of known kinase targets with alignment and structure based approaches to identify and study SDRs. The primary goal of this study was to identify and rationalize SDRs for particular preferences. Importantly, our analysis clearly shows how different positions contribute in unique ways to target site recognition. Many SDR positions were found distal to the substrate binding site. These are harder to rationalize structurally and additional work will need to be done to establish how they relate to target site preference.

The SNF1 mutations of SDRs validated two positions contributing to the expected target preferences - position 164 for the L+4 preference and 189 for the L-5 preference. A recent study also concomitantly predicted position 189 as an L-5 determinant from a comparative structural analysis (Catherine Chen et al. 2017). However, while this residue was mutated and the specificity tested, mutation of 189 always occurred in combination with other kinase residues and so the role of position 189 *per se* as an L-5 SDR remained ambiguous. L+4 specificity to our knowledge was so far uncharacterised and links traditional +1 determinant (position 164) to a distal substrate position (+4).

The study of cancer mutations has revealed that SDRs are commonly mutated as shown previously (Creixell, Schoof, et al. 2015).In addition to previous studies, we observed that SDR mutation burden in cancer can reflect kinase specificities with specific residues being targeted depending on the kinase preferences. Understanding the impact of mutations in kinases will facilitate the classification of cancer mutations into drivers or passenger depending on their functional consequences. Our results suggest that grouping all SDR positions regardless of the kinase specificity will tend to overestimate the impact of mutations since many SDR positions are only relevant for one or few specificities.

The identification of the SDRs allows us to study the evolution of kinase preferences by ancestral sequence reconstruction. The protein kinase domain has been extensively duplicated throughout evolution but very little is known about the process of divergence of kinase target preference. We have shown that kinase orthologs tend to maintain their specificity. This would be expected as they can regulate up to hundreds of targets and a change in specificity would drastically alter the regulation of a large number of proteins. This high conservation of kinase specificity contrasts to the larger divergence rate of kinase target sites (Beltrao et al. 2009; Freschi, Osseni, and Landry 2014; Studer et al. 2016). The evolutionary plasticity of kinase signaling therefore relies primarily on the fast turnover of target sites that can occur without the need for gene duplication.

Examples do still exist however of specificity divergence within kinase families. A previous study has shown how the Ime2 kinases (RCK family) have diverged from the other CMGC kinases in their typical preference for proline at the +1 position (Howard et al. 2014). Here we have traced the putative evolutionary history of the GRK family preference at the -2/-3 positions, which demonstrates divergence of kinase specificity between paralogs and also distantly-related orthologs. An understanding of kinase SDRs will allow for further studies of how the variety of target peptide preferences has come about during evolution and the rate at which kinases can switch their preferences after gene duplication.

Kinase target recognition within the cell is complex and the specificity at the active site is only one of several mechanisms that can determine kinase-substrate interactions (Ubersax and Ferrell 2007). Much additional work is needed to establish a global comprehensive view of kinase target specificity and its evolution.

## Methods

### Kinase specificity models

Known kinase target phosphosites for human, mouse and *S. cerevisiae* were retrieved from HPRD, PhosphoSitePlus, Phospho.ELM and PhosphoGRID (Prasad, Kandasamy, and Pandey 2009; Hornbeck et al. 2015; Dinkel et al. 2011; Sadowski et al. 2013). PhosphoGRID target sites supported exclusively by kinase perturbation followed by MS were excluded and homologous sequences above 85% identity were removed with CD-HIT (Li and Godzik, 2006). Phosphosites mapping to the kinase activation segment were also removed as kinase autophosphorylation sites often conform poorly to kinase consensus motifs (M. L. Miller et al. 2008; Pike et al. 2008). Specificity matrices for each kinase with at least ten target sites were constructed in the form of a position probability matrix (PPM) - 20 x 11 matrices with the columns representing substrate positions -5 to +5; each value representing the empirical residue frequencies for a given amino-acid at a substrate position. For the purpose of scoring only, the PPMs were converted into PWMs by accounting for background amino acid frequencies in the proteome. Cross-validation was used to assess kinase model performance and PWMs with an average area under curve (AUC) value < 0.6 were excluded from further analysis. Too few tyrosine kinase PPMs remained after these filtering steps and we excluded them for further analysis. Kinase group/family/subfamily classifications, were based on the KinBase data resource unless otherwise specified (Manning et al. 2002).

### Position-based clustering of specificity models and sequence alignment-based detection of putative specificity determining residues (SDRs)

Clustering of the PPMs was performed in a position-based manner for each of the five sites N- and C-terminal to the phosphoacceptor amino acid (-5, -4, -3, -2, -1; +1, +2, +3, +4, +5) using the affinity propagation (AP) algorithm (Frey and Dueck 2007) as implemented in the APCluster R package (Bodenhofer, Kothmeier, and Hochreiter 2011). Non-specific clusters or clusters with fewer than 6 constituent kinases were excluded and the clusters were further modified to account for potential false positive and false negative cases (see extended Supplementary Methods).

The MAFFT L-INS-i method was used to generate kinase MSAs for this analysis (Katoh et al. 2005) and the *trim*Al tool was used remove MSA positions containing more than 20 % ‘gap’ sites (Capella-Gutiérrez, Silla-Martínez, and Gabaldón 2009). Kinases were then grouped by specificity according the clustering of their specificity models, as described above, and then iteratively we predicted SDRs for each cluster (e. g. preference for proline at +1 position). To identify putative SDRs, three high-performing methods alignment-based methods were selected (GroupSim, Multi-Relief 3D, SPEER) from previous benchmarking tests (Chakraborty and Chakrabarti 2015). Incorporating predictions from the three methods is expected to achieve higher specificity than any single method (Chakrabarti and Panchenko 2009). A brief explanation of each method is provided in extended Supplementary Methods. As the GroupSim, Multi-Relief 3D, and SPEER methods use distinct schemes for position scoring we selected as putative SDRs those residues lying within a three-way intersection of the top 15 ranked sites for the single methods, as proposed by *Chakrabarti and Panchenko 2009*.

### Kinase-substrate structures

Empirical kinase structures alone and in complex with putative target sequences were retrieved from the PDB (see extended Supplementary Methods). An automated procedure was implemented to identify the kinase substrate-binding residues for the substrate positions -5 to +4 (excluding P0) and all binding residues contacts were categorised as either hydrogen-bonded, ionic, or non-bonded (i.e. hydrophobic or van der Waals). Kinase-substrate homology models were constructed by first superposing the kinase of interest (query) with a template cocrystal structure to achieve a plausible positioning of the substrate peptide. The template kinase were removed and the template peptide mutated *in Silico* (using CHARMM-GUI (Jo et al. 2008)) to the sequence of a known phosphorylation site of the query kinase. After resolving steric clashes between kinase and substrate, the resulting complexes were then subjected to energy minimisation (EM), followed by molecular dynamics (MD) equilibration and production runs using NAMD (Phillips et al. 2005) (see extended Supplementary Methods for additional details).

### Construction of predictive models, cross-validation, and orthology analysis

Naive Bayes (NB) algorithms were used to predict the specificity of protein kinases on the basis of sequence alone. Five separate classifiers were generated, corresponding to the five preferences – P+1, P-2, R-2, R-3, and L-5 – supported by at least 20 kinases. Each classifier was trained on the 119 Ser/Thr kinase sequences of known specificity, where each kinase was labelled (‘positive’ or ‘negative’) according to the clustering of kinase specificity models described above. Leave one-out cross-validation (LOOCV) was then used for each classifier to identify the subset of input SDRs that would optimise the performance of the model on the training data with respect to the AUC. The R libraries klaR and cvTools were used for model generation and cross-validation, respectively (Weihs et al. 2005; Alfons 2012).

For the pan-taxonomic analysis of protein kinase orthologs, orthologous kinase sequences were retrieved automatically from the Ensembl Genomes database (Kersey et al. 2016) using the Ensembl Rest API and were aligned using the MAFFT L-INS-i method. Orthologs were only retrieved for human kinases belonging to the P+1, P-2, R-2, R-3, and L-5 classes (based on naïve Bayes predictions). Kinases within an orthologous group were aligned using the MAFFT L-INS-i method, and residue conservation was assessed on the basis of substitution matrix similarity. Each orthologous sequence was then queried with the specificity model corresponding to the predicted specificity of the human ortholog. Pseudokinases were filtered from the orthologous groups before any analysis was performed.

For the orthology analysis of human, mouse, and yeast kinases, we used the PPMs described above in addition to the 61 yeast specificity matrices presented in (Mok et al. 2010). Before further analysis, the pT and pY sites were removed from each of the peptide screening models, and then the matrices were normalised so that all columns sum to 1. Human and mouse orthologs (if any) for each yeast kinase were then identified using the Ensembl Rest API for the Ensembl Genomes Compara resource (Kersey et al. 2016). The Frobenius distance was calculated then for every possible human-yeast and/or mouse-yeast pair. Distances for PPMs of the same kinase were generated by subsampling phosphorylation sites (n=23) from the same kinase and then calculating all possible pairwise distances between them.

### Analysis of kinase mutations in cancer

Mutation data for primary tumour samples was obtained from The Cancer Genome Atlas (TCGA) (http://cancergenome.nih.gov/). Each Ser/Thr kinase mutation was assigned to the correct protein isoform and then mapped to the corresponding Ser/Thr kinase domain position. All kinase domain positions were categorised as ‘SDR’, ‘Catalytic’, ‘Regulatory’, and ‘Other’. Catalytic and regulatory sites were inferred from the literature. ‘SDR’ sites refers to residues that are both potential SDRs (**Figure 2a**) and often found in close contact with the substrate peptide (**Figure 1b**). ‘Other’ refers to the complement of these three sets relative to the Ser/Thr kinase domain.

### GRK phylogeny and ancestral sequence reconstruction

Protein sequences were retrieved from a taxonomically-broad set of non-redundant proteomes (representative proteomes) (Chuming Chen et al. 2011), and then each representative proteome (rp35) was queried with a hidden Markov model (HMM) of the GRK domain (KinBase) using *HMMsearch* (E = 1e-75) *(Eddy 1998)*. The subfamily classifications of each GRK were then predicted using Kinannote (Goldberg et al. 2013). Sample sequences of the RSK family kinases – the family most similar in sequence to the GRKs – were also included as an expected outgroup in the phylogeny, as were two kinases of the basophilic PKA family. The kinase sequences (GRK kinases plus outgroups) were then aligned using the L-INS-i algorithm of MAFFT (Katoh and Standley 2013), and filtered to remove pseudokinases and redundant sequences (97% threshold), resulting in 163 sequences to be used for phylogenetic reconstruction. A maximum likelihood phylogeny was generated with RAxML using a gamma model to account for the heterogeneity of rates between sites. The optimum substitution matrix (LG) for reconstruction was also determined with RAxML using a likelihood-based approach (Stamatakis 2014). FastML was then used for the ML-based ancestral reconstruction of sequences for all nodes in the phylogeny (Ashkenazy et al. 2012). Sequence probabilities were calculated marginally using a gamma rate model and the LG substitution matrix.

### SNF1 mutant *in vitro* kinase activity assay

The SNF1 plasmid from the Yeast Gal ORF collection was used as a template for directed mutagenesis to create the mutants A218L and V244R. Wild type and mutant plasmids were transformed into a *BY4741 SNF1* KO strain. Cells were grown to exponential phase in SD media lacking uracil, and Snf1 expression was induced with 2% galactose for 8h. Cells were collected by centrifugation at 3200rpm for 5min and kept at -80C. Cell pellets were resuspended in lysis buffer (20mM Tris pH8, 15mM EDTA pH8, 15mM EGTA pH8 and 0.1% Triton X-100) containing a cocktail of protease (cOmplete, from Roche) and phosphates inhibitors (PhosSTOP, from Sigma). Glass beads were added in equal volume (500ml) and cells were lysed by vortexing at 4C. Snf1 immunoprecipitation was performed using rabbit IgG-Protein A agarose beads (Sigma) with rotation for 2h at 4C. Agarose beads were washed 4 times with lysis buffer before mixing with substrates for kinase assay. Kinase assay was performed using AQUA synthetic peptides (Sigma). Each of the 3 kinases was incubated with equal concentration of the 3 synthetic peptides (VQLKRPASVLALNDL, VQDKRPASVLALNDL and VQLKRPASVLAANDL), ATP mix (ATP 300 μM, 15 mM MgCl2, 0.5 mM EGTA, 15 mM β-glycerol phosphate, 0.2 mM sodium orthovanadate, 0.3 mM DTT) and allowed to react for 0, 2, 7 and 20 minutes. The reactions were quenched by transferring the reaction mixture onto dry ice at the corresponding times.

### Mass spectrometry identification and quantification

Kinase reaction products were diluted with 0.1% formic acid in LC-MS grade water and 5 μl of solution (containing 10 pmol of the unmodified peptide substrates) were loaded LC-MS/MS system consisting of a nanoflow ultimate 3000 RSL nano instrument coupled on-line to a Q-Exactive Plus mass spectrometer (Thermo Fisher Scientific). Gradient elution was from 3% to 35% buffer B in 15 min at a flow rate 250 nL/min with buffer A being used to balance the mobile phase (buffer A was 0.1% formic acid in LC-MS grade water and B was 0.1% formic acid in LC-MS grade acetonitrile). The mass spectrometer was controlled by Xcalibur software (version 4.0) and operated in the positive ion mode. The spray voltage was 2 kV and the capillary temperature was set to 255 °C. The Q-Exactive Plus was operated in data dependent mode with one survey MS scan followed by 15 MS/MS scans. The full scans were acquired in the mass analyser at 375-1500m/z with the resolution of 70 000, and the MS/MS scans were obtained with a resolution of 17 500. For quantification of each phosphopeptide and its respective unmodified form, the extracted ion chromatograms were integrated using the theoretical masses of ions using a mass tolerance of 5 ppm. Values of area-under-the-curve were obtained manually in Qual browser of Xcalibur software (version 4.0).

## References

Alfons, Andreas. 2012. “cvTools: Cross-Validation Tools for Regression Models.” R Package Version 0. 3 2 (5).

Amanchy, Ramars, Balamurugan Periaswamy, Suresh Mathivanan, Raghunath Reddy, Sudhir Gopal Tattikota, and Akhilesh Pandey. 2007. “A Curated Compendium of Phosphorylation Motifs.” Nature Biotechnology 25 (3): 285–86.

Ashkenazy, Haim, Osnat Penn, Adi Doron-Faigenboim, Ofir Cohen, Gina Cannarozzi, Oren Zomer, and Tal Pupko. 2012. “FastML: A Web Server for Probabilistic Reconstruction of Ancestral Sequences.” Nucleic Acids Research 40 (Web Server issue): W580–84.

Beltrao, Pedro, Jonathan C. Trinidad, Dorothea Fiedler, Assen Roguev, Wendell A. Lim, Kevan M. Shokat, Alma L. Burlingame, and Nevan J. Krogan. 2009. “Evolution of Phosphoregulation: Comparison of Phosphorylation Patterns across Yeast Species.” PLoS Biology 7 (6): e1000134.

Berthon, Annabel S., Eva Szarek, and Constantine A. Stratakis. 2015. “PRKACA: The Catalytic Subunit of Protein Kinase A and Adrenocortical Tumors.” Frontiers in Cell and Developmental Biology 3 (May): 26.

Biondi, Ricardo M., and Angel R. Nebreda. 2003. “Signalling Specificity of Ser/Thr Protein Kinases through Docking-Site-Mediated Interactions.” Biochemical Journal 372 (Pt 1): 1–13.

Bodenhofer, Ulrich, Andreas Kothmeier, and Sepp Hochreiter. 2011. “APCluster: An R Package for Affinity Propagation Clustering.” Bioinformatics 27 (17): 2463–64.

Brinkworth, Ross I., Robert A. Breinl, and Bostjan Kobe. 2003. “Structural Basis and Prediction of Substrate Specificity in Protein Serine/threonine Kinases.” Proceedings of the National Academy of Sciences of the United States of America 100 (1): 74–79.

Brognard, John, and Tony Hunter. 2011. “Protein Kinase Signaling Networks in Cancer.” Current Opinion in Genetics & Development 21 (1): 4–11.

Capella-Gutièrrez, Salvador, Josè M. Silla-Martínez, and Toni Gabaldón. 2009. “trimAl: A Tool for Automated Alignment Trimming in Large-Scale Phylogenetic Analyses.” Bioinformatics 25 (15): 1972–73.

Chakrabarti, Saikat, and Anna R. Panchenko. 2009. “Ensemble Approach to Predict Specificity Determinants: Benchmarking and Validation.” BMC Bioinformatics 10 (July): 207.

Chakraborty, Abhijit, and Saikat Chakrabarti. 2015. “A Survey on Prediction of Specificity-Determining Sites in Proteins.” Briefings in Bioinformatics 16 (1): 71–88.

Chen, Catherine, Wutigri Nimlamool, Chad J. Miller, Hua Jane Lou, and Benjamin E. Turk. 2017. “Rational Redesign of a Functional Protein Kinase-Substrate Interaction.” ACS Chemical Biology 12 (5): 1194–98.

Chen, Chuming, Darren A. Natale, Robert D. Finn, Hongzhan Huang, Jian Zhang, Cathy H. Wu, and Raja Mazumder. 2011. “Representative Proteomes: A Stable, Scalable and Unbiased Proteome Set for Sequence Analysis and Functional Annotation.” PloS One 6 (4): e18910.

Creixell, Pau, Antonio Palmeri, Chad J. Miller, Hua Jane Lou, Cristina C. Santini, Morten Nielsen, Benjamin E. Turk, and Rune Linding. 2015. “Unmasking Determinants of Specificity in the Human Kinome.” Cell 163 (1): 187–201.

Creixell, Pau, Erwin M. Schoof, Craig D. Simpson, James Longden, Chad J. Miller, Hua Jane Lou, Lara Perryman, et al. 2015. “Kinome-Wide Decoding of Network-Attacking Mutations Rewiring Cancer Signaling.” Cell 163 (1): 202–17.

Dinkel, Holger, Claudia Chica, Allegra Via, Cathryn M. Gould, Lars J. Jensen, Toby J. Gibson, and Francesca Diella. 2011. “Phospho.ELM: A Database of Phosphorylation Sites—update 2011.” Nucleic Acids Research 39 (suppl_1): D261–67.

Doolittle, R. F., D. F. Feng, S. Tsang, G. Cho, and E. Little. 1996. “Determining Divergence Times of the Major Kingdoms of Living Organisms with a Protein Clock.” Science 271 (5248): 470–77.

Eddy, S. R. 1998. “Profile Hidden Markov Models.” Bioinformatics 14 (9): 755–63.

Freschi, Luca, Mazid Osseni, and Christian R. Landry. 2014. “Functional Divergence and Evolutionary Turnover in Mammalian Phosphoproteomes.” PLoS Genetics 10 (1): e1004062.

Frey, Brendan J., and Delbert Dueck. 2007. “Clustering by Passing Messages between Data Points.” Science 315 (5814): 972–76.

Goldberg, Jonathan M., Allison D. Griggs, Janet L. Smith, Brian J. Haas, Jennifer R. Wortman, and Qiandong Zeng. 2013. “Kinannote, a Computer Program to Identify and Classify Members of the Eukaryotic Protein Kinase Superfamily.” Bioinformatics 29 (19): 2387–94.

Holland, P. M., and J. A. Cooper. 1999. “Protein Modification: Docking Sites for Kinases.” Current Biology: CB 9 (9): R329–31.

Hornbeck, Peter V., Bin Zhang, Beth Murray, Jon M. Kornhauser, Vaughan Latham, and Elzbieta Skrzypek. 2015. “PhosphoSitePlus, 2014: Mutations, PTMs and Recalibrations.” Nucleic Acids Research 43 (Database issue): D512–20.

Howard, Conor J., Victor Hanson-Smith, Kristopher J. Kennedy, Chad J. Miller, Hua Jane Lou, Alexander D. Johnson, Benjamin E. Turk, and Liam J. Holt. 2014. “Ancestral Resurrection Reveals Evolutionary Mechanisms of Kinase Plasticity.” eLife 3 (October). https://doi.org/10.7554/eLife.04126.

Jo, Sunhwan, Taehoon Kim, Vidyashankara G. Iyer, and Wonpil Im. 2008. “CHARMM-GUI: A Web-Based Graphical User Interface for CHARMM.” Journal of Computational Chemistry 29 (11): 1859–65.

Kannan, Natarajan, Nina Haste, Susan S. Taylor, and Andrew F. Neuwald. 2007. “The Hallmark of AGC Kinase Functional Divergence Is Its C-Terminal Tail, a Cis-Acting Regulatory Module.” Proceedings of the National Academy of Sciences of the United States of America 104 (4):1272–77.

Kannan, Natarajan, and Andrew F. Neuwald. 2004. “Evolutionary Constraints Associated with Functional Specificity of the CMGC Protein Kinases MAPK, CDK, GSK, SRPK, DYRK, and CK2α.“ Protein Science: A Publication of the Protein Society 13 (8): 2059–77.

Katoh, Kazutaka, Kei-Ichi Kuma, Hiroyuki Toh, and Takashi Miyata. 2005. “MAFFT Version 5: Improvement in Accuracy of Multiple Sequence Alignment.” Nucleic Acids Research 33 (2): 511–18.

Katoh, Kazutaka, and Daron M. Standley. 2013. “MAFFT Multiple Sequence Alignment Software Version 7: Improvements in Performance and Usability.” Molecular Biology and Evolution 30 (4): 772–80.

Kersey, Paul Julian, James E. Allen, Irina Armean, Sanjay Boddu, Bruce J. Bolt, Denise Carvalho-Silva, Mikkel Christensen, et al. 2016. “Ensembl Genomes 2016: More Genomes, More Complexity.” Nucleic Acids Research 44 (D1): D574–80.

Knighton, D. R., J. H. Zheng, L. F. Ten Eyck, V. A. Ashford, N. H. Xuong, S. S. Taylor, and J. M. Sowadski. 1991. “Crystal Structure of the Catalytic Subunit of Cyclic Adenosine Monophosphate-Dependent Protein Kinase.” Science 253 (5018): 407–14.

Kumar, Sudhir, Glen Stecher, Michael Suleski, and S. Blair Hedges. 2017. “TimeTree: A Resource for Timelines, Timetrees, and Divergence Times.” Molecular Biology and Evolution 34 (7): 1812–19.

Lahiry, Piya, Ali Torkamani, Nicholas J. Schork, and Robert A. Hegele. 2010. “Kinase Mutations in Human Disease: Interpreting Genotype-Phenotype Relationships.” Nature Reviews. Genetics 11 (1): 60–74.

Lodowski, David T., Valerie M. Tesmer, Jeffrey L. Benovic, and John J. G. Tesmer. 2006. “The Structure of G Protein-Coupled Receptor Kinase (GRK)-6 Defines a Second Lineage of GRKs.” The Journal of Biological Chemistry 281 (24): 16785–93.

Manning, G., D. B. Whyte, R. Martinez, T. Hunter, and S. Sudarsanam. 2002. “The Protein Kinase Complement of the Human Genome.” Science 298 (5600): 1912–34.

Miller, Chad J., and Benjamin E. Turk. 2018. “Homing in: Mechanisms of Substrate Targeting by Protein Kinases.” Trends in Biochemical Sciences 43 (5): 380–94.

Miller, Martin Lee, Lars Juhl Jensen, Francesca Diella, Claus Jørgensen, Michele Tinti, Lei Li, Marilyn Hsiung, et al. 2008. “Linear Motif Atlas for Phosphorylation-Dependent Signaling.” Science Signaling 1 (35): ra2.

Mok, Janine, Philip M. Kim, Hugo Y. K. Lam, Stacy Piccirillo, Xiuqiong Zhou, Grace R. Jeschke, Douglas L. Sheridan, et al. 2010. “Deciphering Protein Kinase Specificity through Large-Scale Analysis of Yeast Phosphorylation Site Motifs.” Science Signaling 3 (109): ra12.

Mushegian, Arcady, Vsevolod V. Gurevich, and Eugenia V. Gurevich. 2012. “The Origin and Evolution of G Protein-Coupled Receptor Kinases.” PloS One 7 (3): e33806.

Ochoa, David, David Bradley, and Pedro Beltrao. 2018. “Evolution, Dynamics and Dysregulation of Kinase Signalling.” Current Opinion in Structural Biology 48 (February): 133–40.

Onorato, J. J., K. Palczewski, J. W. Regan, M. G. Caron, R. J. Lefkowitz, and J. L. Benovic. 1991. “Role of Acidic Amino Acids in Peptide Substrates of the Beta-Adrenergic Receptor Kinase and Rhodopsin Kinase.” Biochemistry 30 (21): 5118–25.

Pearson, R. B., and B. E. Kemp. 1991. “Protein Kinase Phosphorylation Site Sequences and Consensus Specificity Motifs: Tabulations.” Methods in Enzymology 200: 62–81.

Phillips, James C., Rosemary Braun, Wei Wang, James Gumbart, Emad Tajkhorshid, Elizabeth Villa, Christophe Chipot, Robert D. Skeel, Laxmikant Kalè, and Klaus Schulten. 2005. “Scalable Molecular Dynamics with NAMD.” Journal of Computational Chemistry 26 (16): 1781–1802.

Pike, Ashley C. W., Peter Rellos, Frank H. Niesen, Andrew Turnbull, Antony W. Oliver, Sirlester A. Parker, Benjamin E. Turk, Laurence H. Pearl, and Stefan Knapp. 2008. “Activation Segment Dimerization: A Mechanism for Kinase Autophosphorylation of NonL Jconsensus Sites.” The EMBO Journal 27 (4): 704–14.

Pinna, L. A., and M. Ruzzene. 1996. “How Do Protein Kinases Recognize Their Substrates?” Biochimica et Biophysica Acta 1314 (3): 191–225.

Prasad, T. S. Keshava, Kumaran Kandasamy, and Akhilesh Pandey. 2009. “Human Protein Reference Database and Human Proteinpedia as Discovery Tools for Systems Biology.” Methods in Molecular Biology 577: 67–79.

Sadowski, Ivan, Bobby-Joe Breitkreutz, Chris Stark, Ting-Cheng Su, Matthew Dahabieh, Sheetal Raithatha, Wendy Bernhard, et al. 2013. “The PhosphoGRID Saccharomyces Cerevisiae Protein Phosphorylation Site Database: Version 2.0 Update.” Database: The Journal of Biological Databases and Curation 2013 (May): bat026.

Saunders, Neil F. W., Ross I. Brinkworth, Thomas Huber, Bruce E. Kemp, and Bostjan Kobe. 2008. “Predikin and PredikinDB: A Computational Framework for the Prediction of Protein Kinase Peptide Specificity and an Associated Database of Phosphorylation Sites.” BMC Bioinformatics 9 (1): 245.

Stamatakis, Alexandros. 2014. “RAxML Version 8: A Tool for Phylogenetic Analysis and Post-Analysis of Large Phylogenies.” Bioinformatics 30 (9): 1312–13.

Stenberg, K. A., P. T. Riikonen, and M. Vihinen. 2000. “KinMutBase, a Database of Human Disease-Causing Protein Kinase Mutations.” Nucleic Acids Research 28 (1): 369–71.

Studer, Romain A., Ricard A. Rodriguez-Mias, Kelsey M. Haas, Joanne I. Hsu, Cristina Vièitez, Carme Solè, Danielle L. Swaney, et al. 2016. “Evolution of Protein Phosphorylation across 18 Fungal Species.” Science 354 (6309): 229–32.

Ubersax, Jeffrey A., and James E. Ferrell Jr. 2007. “Mechanisms of Specificity in Protein Phosphorylation.” Nature Reviews. Molecular Cell Biology 8 (7): 530–41.

Varjosalo, Markku, Salla Keskitalo, Audrey Van Drogen, Helka Nurkkala, Anton Vichalkovski, Ruedi Aebersold, and Matthias Gstaiger. 2013. “The Protein Interaction Landscape of the Human CMGC Kinase Group.” Cell Reports 3 (4): 1306–20.

Weihs, Claus, Uwe Ligges, Karsten Luebke, and Nils Raabe. 2005. “klaR Analyzing German Business Cycles.” In Data Analysis and Decision Support, 335–43. Studies in Classification, Data Analysis, and Knowledge Organization. Springer, Berlin, Heidelberg.

Xu, Qifang, Kimberly L. Malecka, Lauren Fink, E. Joseph Jordan, Erin Duffy, Samuel Kolander, Jeffrey R. Peterson, and Roland L. Dunbrack Jr. 2015. “Identifying Three-Dimensional Structures of Autophosphorylation Complexes in Crystals of Protein Kinases.” Science Signaling 8 (405): rs13.

Zhu, Guozhi, Koichi Fujii, Natalya Belkina, Yin Liu, Michael James, Juan Herrero, and Stephen Shaw. 2005. “Exceptional Disfavor for Proline at the P+ 1 Position among AGC and CAMK Kinases Establishes Reciprocal Specificity between Them and the Proline-Directed Kinases.” The Journal of Biological Chemistry 280 (11): 10743–48.

